# An analytical framework for interpretable and generalizable ‘quasilinear’ single-cell data analysis

**DOI:** 10.1101/2020.04.12.022806

**Authors:** Jian Zhou, Olga G. Troyanskaya

**Affiliations:** Lyda Hill Department of Bioinformatics, University of Texas Southwestern Medical Center, Texas, United States of America; Lewis-Sigler Institute for Integrative Genomics, Princeton University, Princeton, New Jersey, United States of America; Flatiron Institute, Simons Foundation, New York, New York, United States of America; Department of Computer Science, Princeton University, Princeton, New Jersey, United States of America

## Abstract

Scaling single-cell data exploratory analysis with the rapidly growing diversity and quantity of single-cell omics datasets demands more interpretable and robust data representation that is generalizable across datasets. To address this challenge, here we developed a novel ‘quasilinear’ framework that combines the interpretability and transferability of linear methods with the representational power of nonlinear methods. Within this framework, we introduce a data representation and visualization method, GraphDR, and a structure discovery method, StructDR, that unifies cluster, trajectory, and surface estimation and allows their confidence set inference. We applied both methods to diverse single-cell RNA-seq datasets from whole embryos and tissues. Unlike PCA and t-SNE, GraphDR and StructDR generated representations that both distinguished highly specific cell types and were comparable across datasets. In addition, GraphDR is at least an order of magnitude faster than commonly used nonlinear methods. Our visualizations of scRNA-seq data from developing zebrafish and Xenopus embryos revealed extruding branches of lineages from a continuum of cell states, suggesting that the current branch view of cell specification may be oversimplified. Moreover, StructDR identified a novel neuronal population using scRNA-seq data from mouse hippocampus. An open-source python library and a user-friendly graphical interface for 3D data visualization and analysis with these methods are available at https://github.com/jzthree/quasildr.

## Introduction

Single-cell gene expression^1–3^ and chromatin profiling techniques^4–6^ have vastly expanded our understanding of cell state variation and heterogeneity. For instance, single-cell RNA-seq identified a novel, rare cell type in lungs, the CFTR-expressing pulmonary ionocyte, that is likely critical to cystic fibrosis pathology^7,8^. Moreover, the quantity and scale of single-cell datasets are rapidly increasing, which provide groundbreaking potential of new discoveries for key cell types and cellular states that underlie specific biological and clinical conditions through integrating knowledge across datasets. However, integrative analyses of single-cell datasets are challenging for a variety of reasons, including characteristics of single-cell omics data such as very high number of cells, biological complexity and heterogeneity of samples, and very noisy measurements, and a lack of data representation methods that address while being genearalizable across datasets.

Single-cell exploratory analysis methods, including visualization methods and approaches for trajectory estimation, rely on either linear or non-linear data representation, each of which presents important limitations to the single-cell data analysis. Linear dimensionality reduction methods, including Principal Component Analysis (PCA) and Independent Component Analysis (ICA), provide clear interpretation via linear maps with uniform interpretation of directions and distances in the representation space. Furthermore, it is simple to apply the same low dimensional projection to different datasets, producing comparable representations. However, linear dimensionality reduction methods typically cannot efficiently represent cell identities in single-cell data: spatial adjacency in low-dimensional representations is not as good predictor of similarity in overall expression state compared to nonlinear methods.

These limitations have led to wide use of nonlinear representations, such as as t-distributed stochastic neighbor embedding (t-SNE) or UMAP, but they generally lack many of the desirable properties enjoyed by linear methods such as interpretability and comparability. For example, these representations are not comparable across datasets, limiting analysis such as analyzing shared cell types across tissues. Similarly, for trajectory estimation, trajectories are essentially specialized nonlinear representations of the data, and the lack of analytical tractability from existing methods prevents applying statistical inference to analyze uncertainties of the resulting trajectories. This makes drawing robust conclusions difficult when, for example, attempting to distinguish highly similar lineages,

These limitations present a practical barrier to compare or integrate datasets at scale, Yet, interpretable and comparable data representations are essential for the analysis of, for example, multiple disease conditions for the same tissue or different regions within a given organ, where directly comparable and interpretable representations will be crucial, if not essential. In addition, the quest to identify increasingly specialized cell types, cell states, and trajectories at scale demands a principled statistical approach to distinguish signal from noise, such as inference of confidence sets for the extracted structures.

We hypothesize that the difficulty of linear dimensionality reduction for single-cell data arises from the high level of noise or stochasticity: high dimensionality is necessary to capture similarities between cells, and this renders low dimensional linear representations less appealing. Indeed, all popular nonlinear methods for single-cell omics data use high-dimensional information, which is often represented by distances between cells from high dimensional input, thus effectively reducing the effect of noise. We reasoned that allowing information-sharing across cells leveraging high-dimensional information could improve the quality of cell state representation while preserving the linear space and its interpretability.

We therefore developed a novel ‘quasilinear’ framework for exploratory analysis of single-cell omics data, which includes both visualization and structure extraction such as trajectory estimation. We define quasilinear representations as a special group of nonlinear representations that exactly or approximately preserve the interpretability of a linear subspace but are better than linear methods with respect to cell state representation quality or other desired properties. Each quasilinear representation is strongly connected to a linear representation in an analytically tractable manner. In effect, this quasilinear framework combines the advantages of both linear and non-linear methods. We developed two distinct quasilinear methods that complement each other: an interpretable and transferable dimensionality reduction and visualization method, GraphDR, and a general structure extraction method, StructDR, that unifies cluster, trajectory, and surface estimation under the same framework and enables inference of confidence sets for these structures. An open-source python library and an user-friendly, interactive graphical interface are provided for interactive analysis and visualization.

## Results

### Quasilinear dimensionality reduction for visualizing single-cell data: GraphDR

To overcome the limitations of linear representations in single-cell data while preserving their benefits, we developed GraphDR, a graph-based quasilinear data representation and visualization method that addresses the limitations of linear representations in single-cell data while preserving its benefits (**Figure 1a**). We achieved these desired properties by considering a flexible class of ‘quasilinear’ methods. This class of quasilinear transform improves over linear methods but maintains interpretability by introducing nonlinearity specifically for information sharing across cells. Briefly, quasilinear methods apply: (1) a feature (e.g. gene) space transformation *W*, as in linear methods, and (2) an interpretability-preserving cell space transformation *K* that introduces nonlinearity and improves cell state representation (**Methods**).

**Figure 1.**
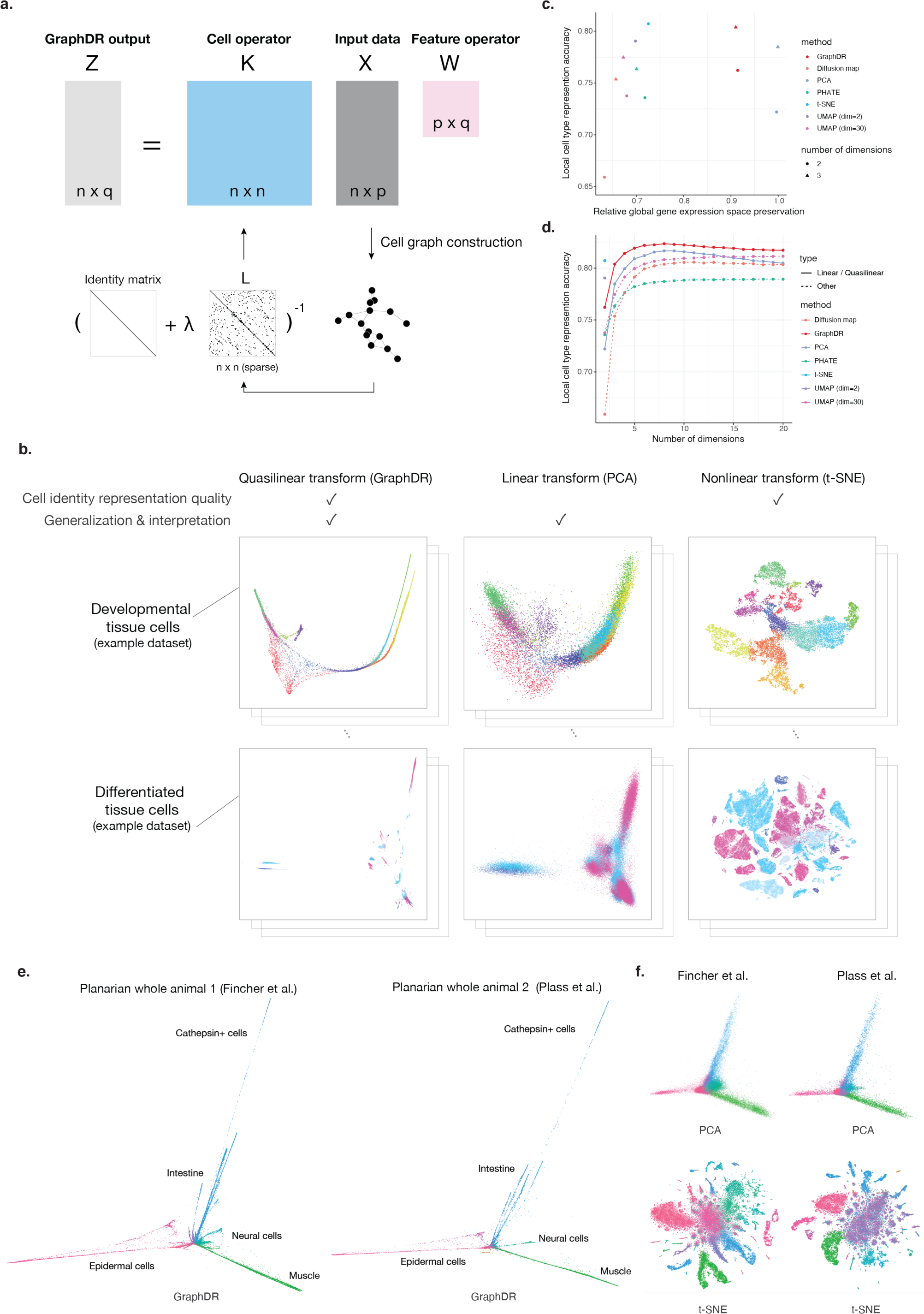
A quasilinear transform method that captures the structure of single-cell data while preserving interpretability and transferability. **a**. Schematic overview of the quasilinear transform method GraphDR for single-cell omics data representation and visualization. GraphDR approximately preserves the structure and interpretability of a corresponding linear transform. **b**. Visualization of two example datasets of developmental trajectory^9^ (top) and mature cell types^11^ (bottom) using GraphDR and representative linear and nonlinear methods, PCA and t-SNE. GraphDR is applied without rotation relative to PCA (Methods). **c**. Comparison of single-cell data dimensionality reduction methods in representing cell type identity and preserving gene expression space. Y-axis shows the accuracy of recovering cell type information from its nearest neighbor in the representation. X-axis shows preservation of the input linear space measured in correlation of pairwise distance. Both two-dimensional (triangles) and three-dimensional (solid dot) representations are compared. **d**. Cell type identity representation accuracies in multiple numbers of dimensions for single-cell data dimensionality reduction methods. **e-f**. Quasilinear transform facilitates comparison across datasets, balancing advantages of linear and nonlinear transform. Two planarian single-cell datasets (e. left panel and f. top panel: Fincher et al. 2018; e. right panel and f. bottom panel: Plass et al. 2018) were processed with a representative linear transform PCA, a nonlinear transform t-SNE, and quasilinear method GraphDR.

In particular, GraphDR applies a cell space transformation derived from the analytical solution of a graph-based optimization problem that provide information sharing across cells connected in a graph (**Figure 1a, Methods**). The graph can be constructed with cell state similarities in high-dimensional input data and incorporate experimental designs when appropriate. The optimization problem then simutaenously optimizes the reconstruction of the input data and the consistency with the graph. The existence of a closed-form solution also makes GraphDR analytically tractable and allows ultrafast computation.

As a proof-of-principle, we first applied GraphDR, PCA, and t-SNE to two distinct single-cell RNA-seq datasets, representing a developing trajectory of mouse hippocampus cell types^9^ and mature mouse brain cell types^10^ (**Figure 1a**). GraphDR generated representations that preserved the interpretability of subspace like PCA and resolved the different cell types like t-SNE (**Figure 1b-d**). Therefore, importantly, this gain of interpretability was achieved without a loss of accuracy. Moreover, in a large-scale quantitative benchmark across a diverse set of seven single-cell datasets, GraphDR distinguished cell types/states as well as several current state-of-the-art nonlinear methods, measured by consistency of nearest neighbors in dimensionality-reduced embedding with literature-based cell type identities (**Figure 1c-d, Methods**).

GraphDR also facilitates direct comparisons across datasets. To demonstrate, we used GraphDR to analyze two planarian *Schmidtea mediterranea* whole-animal single-cell RNA-seq datasets by two different labs^12,13^. GraphDR generated representations that could both distinguish all cell types and be compared across datasets (**Figure 1e**). In contrast linear PCA representations were comparable but did not resolve specific cell types, whereas nonlinear t-SNE representations resolved cell types but were not comparable across datasets (**Figure 1f**). Notably, no special dataset alignment is typically necessary to generate comparable representations for visual comparisons, but GraphDR is also uniquely capable of applying dataset alignment or batch correction in the representation computation procedure for more precise, fine-grained comparison of datasets (**Supplementary Figure 1-2**).

### Extensibility and scalability of GraphDR

GraphDR can also incorporate experimental design information into the analysis whenever appropriate. For example, we can incoportate temporal and batch information by encoding them in the graph construction step of GraphDR. Specifically, we connect nearest neighbor cells between adjacent time points or between two different batches at the same time point (**Supplementary Figure 2, Methods**).

To illustrate this, we applied GraphDR to single-cell datasets that characterize complex developmental progression landscapes. We visualized single-cell RNA-seq datasets from developing zebrafish embryos scRNA-seq dataset (time-series design; **Supplementary Figure 3**) and Xenopus embryos (batch + time-series design; **Supplementary Figure 4**). Incorporating design information was critical for correctly representing the developmental progression correctly. Interestingly, the visualization of each developmental landscape revealed extruding branches of lineages from a continuum cell states (**Supplementary Figure 3** and **Supplementary Figure 4**). These data suggest that the current branch view of cell fate specification is an oversimplification, and that a more sophisticated paradigm is needed.

Single-cell dataset size are growing in size, so designing fast algorithms that scale with dataset size is essential. GraphDR takes 5 minutes to analyze a large 1.3 million cells on a typical modern server machine (2x Xeon Gold 6148), which is 10x faster than UMAP (52 minutes), currently one of the fastest nonlinear dimensionality reduction methods.

For very large datasets that will become available in the near future, we developed a GPU-accelerated version of GraphDR, which takes only 1.5 minutes to analyze 1.3 million cells dataset and 18 minutes for 10 million cells simulated dataset (1x Tesla V100). To achieve this performance, we optimized each major step of computation to use fast algorithms and implementations. The two major steps in the GraphDR algorithms are graph construction and solving the output Z. To scale the graph construction step, we leveraged recent progress in fast approximate KNN algorithms such as hierarchical navigable small-world graphs (HNSW)^14^. To compute output *Z*, we avoided explicit computation of *K*, but instead solved *Z* with a linear solver. To solve the linear systems efficiently with modern multicore architecture, we used libraries with highly optimized linear algebra routines, including taking advantage of CUDA-based GPU computation.

### A unified framework for single-cell cluster, trajectory, and surface structure discovery: StructDR

Visualization methods provide an intuitive and flexible representation of the structure of the data. However, quantitatively defined structures such as clusters and trajectories often need to be extracted in order to perform the detailed analysis of cell types, cell states, and developmental trajectories. For instance, identifying genes differentially expressed between cell types or along a differentiation trajectory requires extraction of cluster or trajectory structures. Despite significant advances, existing methods are limited in the complexity of structures that they can represent; for example, no approach for unsupervised surface or mixed-dimensional structure discovery exists. Furthermore, current methods do not allow statistical inference of uncertainties, which is essential for assessing the robustness of conclusions.

We developed a quasilinear approach, StructDR, that leverages the nonparametric density ridge estimation (NRE) method^16–18^. It unifies the estimation of single-cell clusters and trajectories, with new, complex structure types such as surfaces and allowed rigorous estimation of statistical confidence of these structures via bootstrapping (**Figure 2a, Methods**). We found that StructDR provided superior performance on trajectory estimation evaluated on a diverse collection of single-cell RNA-seq datasets (**Figure 2b**). Moreover, the richer structural representation capability offered by StructDR enabled capturing complex heterogeneities, such as differentiating cells in different cell cycle stages of the cell types (**Figure 2c**).

**Figure 2.**
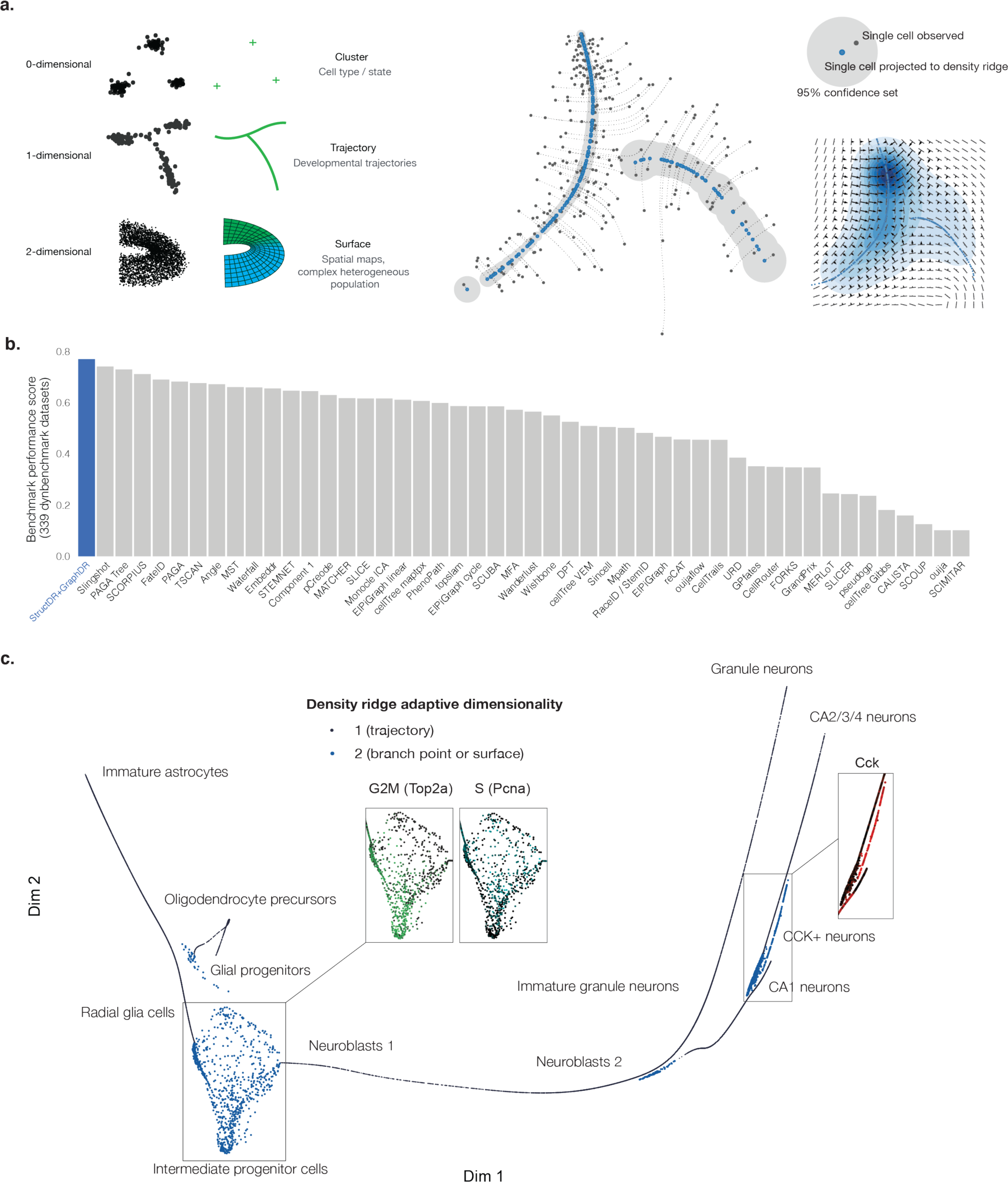
Density-based generalized trajectory estimation and inference. **a**. Schematic overview of the StructDR framework. Left panel: zero-, one-, and two-dimensional density ridges and examples of corresponding biological structures. Mid panel: an example of trajectory estimation (1-dimensional density ridge) based on myoblast single-cell RNA-seq data^20^. The original cell positions are shown in black dots; the projected positions are shown in blue; and the projection lines are shown in dotted lines. Gray shades show confidence sets of trajectory positions. Right panel: the top plot shows an annotated example of confidence set estimation. The bottom plot depicts the elements of the subspace constrained mean-shift algorithm^17^ for performing nonparametric ridge estimation; the arrows indicate gradient vectors of the probability density function; the bars indicate the directions of first eigenvectors of the Hessians of the log probability density function; the kernel density estimator-based density function is shown with the contour plot; the estimated trajectory positions are shown in blue dots. **b**. Performance of StructDR+GraphDR tested on a published large-scale benchmark of 339 datasets. The performance scores are computed based on Saelens et al. 2019^19^. **c**. Trajectory identification with adaptive dimensionality example on a hippocampus developmental trajectory single-cell dataset^9^.

NRE does not make any strong assumptions about the structure of the data, such as the existence of a hidden manifold, in contrast to many existing methods. Instead the problem of identifying structures is cast as discovery of k-dimensional density ridges. Structures such as clusters, trajectories, and surfaces correspond to zero-, one-, and two-dimensional density ridges, which are uniquely defined given any smooth density function of cells estimated from single-cell data. A key additional advantage of this quasilinear approach is that the estimated trajectory positions are directly interpretable as cell states in the input space, and, conversely, all cell states, including cells not observed in the input set, can be mapped to positions on density ridges (**Methods**).

The original NRE method becomes less statistically efficient with higher dimensional, single-cell data (e.g. when the number of principal components used > 6), limiting the method from utilizing all available information. To address this challenge, we utilize GraphDR to generate a quasilinear representation of the data, enabling StructDR to use of all informative principal components for improved structure estimation.

Importantly, we show that StructDR’s gain in representation capabilty does not compromise in accuracy. To evaluate the performance of our framework, we benchmarked the trajectory estimation performance with a large benchmark dataset created by Saelens et al^19^. This dataset includes 339 diverse real and synthetic single-cell datasets. Our trajectory estimation framework showed top performance across all datasets in this benchmark compendium (**Figure 2b, Supplementary Figure 5**).

A key benefit of StructDR is that it captures the full complexity of single-cell data in complex structures, including zero, one, two-dimensional, mixed-dimensional representations (**Figure 2a**,**c, Supplementary Figure 6**). For example, we found that the transcriptomic states of cells at branch points or of differentiating cells going through the cell cycle vary in more than one direction, making clusters of one-dimensional trajectories insufficient for representing the molecular state variations that underlie this cell heterogeneity(**Figure 2c, Supplementary Figure 6**).

Furthermore, our method can adaptively select dimensionality for representing each cell (**Methods**). We used StructDR to analyze scRNA-seq data of hippocampus cell types in perinatal, juvenile, and adult mice. StructDR captured the cellular heterogeneity of neuronal progenitor cells going through the cell cycle by a two-dimensional surface instead of arbitrarily mapping these cells to one-dimensional trajectories (**Figure 2c**). Furthermore, our framework identified a novel CCK+ neurons population between CA1 and CA2/3/4 branches within the hippocampus (**Figure 2c**). This population was apparent with the adaptive dimensionality structure representation, and was not reported in the previous analysis of this dataset with standard methods^9^.

### Statistical inference of uncertainties in trajectory estimation

As the growing quantity of single-cell data empowers our ability to uncover even subtle differences between cells, it is increasingly important to reliably distinguish signal from noise in an automated and scalable manner. This distinction requires a robust statistical characterization of structural representations generated by single-cell analysis methods. Unlike prior single-cell trajectory estimators, NRE is uniquely capable of estimating confidence sets of ridge positions when applied with a linear representation (nonlinear or quasilinear representations are not yet supported in theoretical results)^18^ (**Figure 2a**).

To demonstrate that NRE-based confidence sets effectively controlled coverage probability of ground-truth trajectory positions in trajectory estimation, we performed real data-based simulations (**Methods**). We found that the confidence interval coverage probabilities are accurately controlled (**Supplementary Figure 7**) as expected from theoretical results.

### Interactive 3D interface for single-cell data visualization exploratory analysis

Interactivity is a cornerstone for exploratory data analysis. To facilitate interaction and to make our tools accessible to a broad range of researchers, we developed an interactive analysis and visualization interface, Trenti, as part of an open source python package, Quasildr (https://github.com/jzthree/quasildr), which implements GraphDR and StructDR methods as described above. Trenti is a feature-rich single-cell omics data exploratory analysis and visualization interface (**Figure 3**). In addition to enabling all of the analyses that we presented so far, we included additional software features, such as the integration of popular dimensionality reduction and clustering methods, built-in gene expression explorer to perform flexible gene expression queries, interactive selector for genes and cells, and flexible visualization adjustments by the user. The interface features an interactive three-dimensional visualization, which we demonstrate has a clear measurable advantage over 2D in many scenarios (**Figure 1d**).

**Figure 3.**
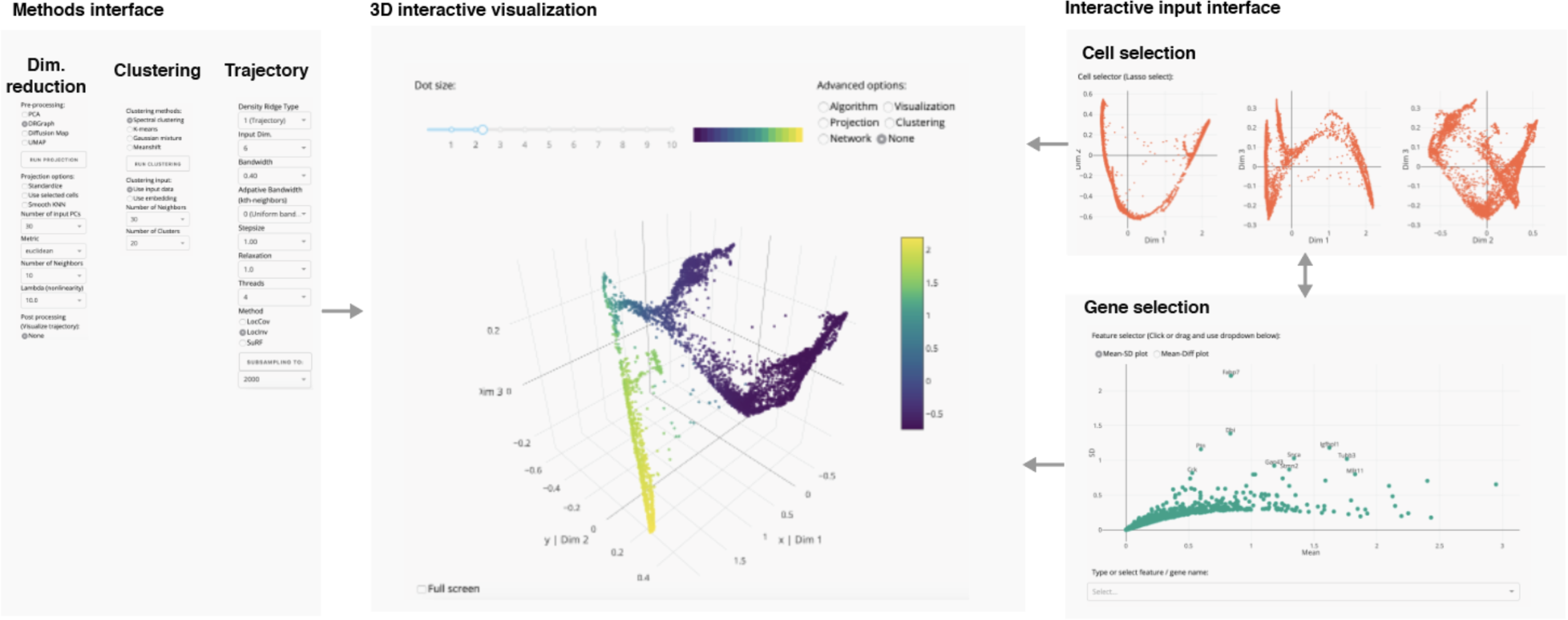
Graphical interface for interactive single-cell visualization and analysis. The elements of the interface include a method interface for different types of analyses - dimensionality reduction, clustering, and trajectory analysis (left), a 3D interactive cell visualization interface (mid), and an interactive filter interface including cell selection and gene selection tools (right). All interfaces are updated upon receiving any input.

## Discussion

Our work presented a quasilinear single-cell data analysis framework that facilitates dataset comparison and integration by providing interpretable representations that are easily transferable. NRE^18^ allows estimating statistical confidence sets of single-cell data density ridge positions, but further work may provide even more flexible and powerful inference. For example, only linear representations such as those provided by PCA are currently well-supported by statistical theory, as most nonlinear representations including quasilinear representations introduce dependencies that complicate theoretical guarantees for bootstrap. In addition, other important properties of interest, such directionality (as opposed to just positions) of a trajectory, have no known methods for confidence set inference.

The scope of “quasilinear” methods is much broader than the algorithms that we discussed above, opening the door for designing quasilinear methods with other desired properties. These methods are also potentially applicable to visualization and exploratory analysis of other high-dimensional data beyond single-cell data applications.

## Methods

### GraphDR - a quasilinear data representation method

We propose a class of quasilinear dimensionality reduction methods, which are nonlinear methods that produce representations that aims to maximally preserve the interpretability of a corresponding linear subspace, while allowing other desired properties unachievable by linear method such as information sharing across cells. We believe it is beneficial to provide a unified viewpoint for this class of methods sharing the same properties.

To design a quasilinear representation method, we first propose the form *Z* = *KXW*, analogous to the linear dimensionality reduction *Z* = *XW*, to allow fast computation and analytical tractability. *Z* represents the data representation output matrix (*n* × *d*, where *n* is the number of samples and *d* is the number of output dimensions), *X* is the input data (*n* × *c*, where *n* is the number of samples and *c* is the number of input dimensions). *W* and *K* are matrices that apply feature (e.g. gene) space and cell space linear transformations that are of shape *d* × *c* and *n* × *n* respectively. In other words, we apply both a linear projection on feature space *W* like linear methods, and an additional linear transform on cell space *K* which is also derived from *X*. The addition of cell space operator *K* allows much greater flexibility in the transformation, which can be exploited to improve the quality of the representation. For example, setting *K* to a block-diagonal matrix with all entries within a block equal to 1/block-size can move all cells within one block to their average position, leading to clustering-like behavior. In theory, an ideal *K* can move all cells within the same ground truth state to the same position asymptotically in the limit of large number of cells.

For the design of K in GraphDR, we use *K* = (*I* + λ*L*)^−1^, which is motivated by the solution to loss function

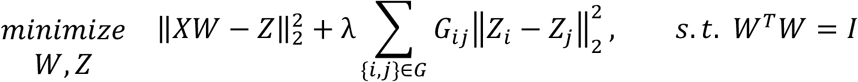

where the first term is the typical PCA loss and the second term is a graph-based regularization term that encourages cells connected in the graph to be close to each other. *L* is the graph Laplacian matrix of graph *G*. The second loss term is also shared by a related nonlinear representation method Laplacian eigenmap. Compared to Laplacian eigenmap, it allows a quasilinear interpretation not available to Laplacian eigenmap and avoids the difficulty when the graph contains disconnected components. The analytical solution to the optimization problem is *Z* = (*I* + λ*L*)^−1^*XW*, where *W* is the top-n eigenvectors of *X*^*T*^(*I* + λ*L*)^−1^*X* where *n* is the dimensionality of *Z*. The existence of an analytical solution makes it much easier to be analyzed, modified, and incorporated in downstream analyses compared to methods that do not.

For graph *G*, a practical and empirically well-performing choice for GraphDR is the nearest-neighbors graph. The graph construction process can also incorporate experimental design or prior knowledge information. For example, nearest neighbors or mutual nearest neighbours between batches can be connected to address batch effect during computation of representation, or nearest neighbors between consecutive time points can be connected when temporal information is available.

GraphDR can also be applied with a predefined *W* matrix or without reducing the dimensionality. Preserving the input data dimensionality is useful for preserving the ability of choosing a linear subspace to visualize after applying the transformation, allowing more flexible comparison of datasets processed separately. With *W* fixed to be identity matrix in order to preserve the original input space, the problem becomes

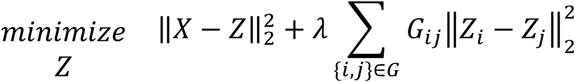

The solution is also simplified as *Z* = *KX*, while K remains unchanged as (*I* + λ*L*)^−1^.

### Computational efficiency optimization

With the constant growth in single-cell dataset size, it is extremely important to design fast algorithms that scale with the dataset size. We have optimized the performance of GraphDR, resulting in an ultrafast method that takes only 1.5 min for 1.3 million cell datasets. To achieve this, each major step of computation has been optimized to use fast algorithms and implementations.

The two major steps in the GraphDR algorithms are graph construction and the final step of solving the output Z. To scale the graph construction step, we leveraged recent progresses in fast approximate KNN algorithms (ANN). Exact KNN algorithms based on ball-tree or KD-tree fit the need for small to medium-sized datasets but do not scale to very large number of cells. For ANN algorithms, we support both the HNSW method^14^ from NMSlib written in C++ with python binding, and a pure python implementation of NN-descent method^21^ built into the package (originally implemented by the UMAP package^22^). The HNSW option is faster and used for our performance test.

In the final step of computing *Z*, for problems with a large number of cells, it is much faster to avoid explicit computation of *K* but solving *Z* with a linear solver. This is because the inverse of *K, I* + λ*G* is sparse and thus allows fast computation. To implement the linear solver efficiently with modern multicore architecture we used libraries with highly optimized linear algebra routines, including taking advantage of CUDA-based GPU computation which gives the best performance.

### Single-cell data structure discovery with nonparametric density ridge estimation

We propose to unify cluster, trajectory, and surface estimation by formulating it as a nonparametric density ridge estimation problem. The nonparametric density ridge estimation problem can be solved via the subspace constrained mean shift (SCMS) algorithm^17,23^. The statistical theory of nonparametric density ridge estimation is described in detail in ^16,18^.

Briefly, density ridge generalizes the concept of local maxima in probability density functions, whereas local maxima correspond to zero-dimensional ridges, trajectories correspond to one-dimensional ridges, and surfaces correspond to two-dimensional ridges. Therefore, zero-dimensional ridge estimation is equivalent to clustering with mean-shift algorithm, and one- and two-dimensional ridge estimation corresponds to trajectory and surface identification algorithms. In addition, the algorithm can project all cells, including unobserved cell states, to their corresponding positions on the density ridges.

In a N-dimensional space and positions of cells in this space representing cell states from single-cell data, nonparametric ridge estimation identifies positions that satisfy the condition *R* = {*x*: ∥*G*_*d*_ (*x*)∥ = 0, λ_,*d*+1_(*x*) < 0}, where *d* is the dimensionality of the density ridge^16^. ∥*G*(*x*)∥ = 0 is the key condition that the projected gradient of the probability density function at this position equals zero, which we will explain in more detail below. λ_*d*+1_(*x*) < 0 is a stability condition that requires trajectory to include points which are ‘local maxima’ instead of ‘local minima’ in probabilistic density function, where λ_*d*+1_(*x*) is the *d* + 1 th largest eigenvalue of the Hessian matrix of probability density function. The projected gradient is computed by projecting out the gradient in the subspace spanned by the top-d eigenvectors. Kernel density estimator with Gaussian kernel is used to estimate the probability density function as it provides a smooth density function that allows fast computation of derivatives.

Nonparametric ridge estimation with ridge dimensionality being one can be considered a principal curve method^17^. Comparing to other principal curve approaches, it has the advantage that the density ridge is uniquely defined and can be efficiently identified using the methods described below once the probability distribution is estimated. We use the subspace-constrained mean-shift (SCMS) algorithm to simultaneously solve the problem of identifying the ridges and projecting individual cells to the trajectory.

Briefly, the algorithm iteratively moves any point toward the projected gradient direction of the density function, until it converges to a point at which the gradient direction point to the direction that is projected out, which is in this case the first eigenvector of the hessian of the log density function. To adaptively decide the step size of each update along the projected gradient direction is derived from mean-shift algorithm, which can be derived from fixed-point iteration and has good convergence properties. A smaller step size than the mean shift step size can be chosen to more accurately integrate over the projected gradient curve and thus projecting single-cells more accurately.

As described in Chen et al.^18^, the bootstrap confidence set can be constructed through the following procedure: 1. first generate *N* bootstrap samples by via sampling with replacement; then estimate density ridges for each bootstrap sample; 2. for each position in the density ridge set estimated from the original sample, calculate the distance to its nearest position in each bootstrap sample density ridge set; 3. take *α* -upper quantile of the distances *t*_α_, and the *α* -confidence set of each estimated ridge position is constructed as a sphere of radius *t* _α_ centered at the estimated ridge position.

Two properties of this approach of constructing confidence sets for density ridge positions should be noted. First is that the definition of true density ridges considered are the density ridges of the smoothed true distribution when applied the same KDE kernel. Second is that the theoretical asymptotic properties of bootstrap statistical inference is only proved for linear representations.

### Adaptive density ridge dimensionality

To allow flexible representations of data containing complex structure with different dimensionalities, we propose the use mixed-dimensionality representation that adaptively determines ridge dimensionality. Empirically, we find mixed one and two-dimensional density ridges to be a robust and informative representation of data structure, which can be used in conjunction with zero-dimensional density ridges. Specifically, we modified the SCMS method to adaptively determine, for any position in the space, the ridge dimensionality between d=1 (trajectory) mode vs d=2 (branch point or surface) mode at every iteration. This decision is based on the eigengap between the first and second eigenvalues of the Hessian matrix of the log probability density function. Specifically if 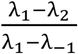 surpassed the specified threshold d=1 is used, otherwise d=2 is used.

### Simulation study for evaluation of confidence set

We used inferred trajectory from a real dataset^9^ as the ground truth to generate synthetic datasets. 100 simulated datasets are generated by adding independent Gaussian noise to the samples from ground truth trajectory density function. For each simulated dataset, 20 bootstraps were used to construct confidence set of trajectory positions based on distance from bootstrapped trajectories to the estimated trajectory as described in ^18^. The estimated confidence sets with a coverage probability were then compared with the ground truth to decide the true proportion of times any point at the ground truth trajectory is covered by the confidence set constructed.

### Graph construction from density ridge positions

In StructDR output, density ridges are represented by positions of data points projected to the these ridges. To allow more flexible applications we construct graph representations from these projected points. To do so, we first construct a candidate graph connecting k-nearest neighbors in both the projected cell space and in the input cell space. The candidate graph is then simplified by choosing only the one nearest neighbors in 2^*d*^ orthogonal or opposite directions in the projected cell space, where *d* is the density ridge dimensionality (e.g. two nearest neighbors are chosen for *d* = 1 or trajectories, and 4 are chosen for *d* = 2 or surfaces). We chose the directions based on first-d eigenvectors of the Hessian, leveraging the observation that density ridges typically extend on same directions as these eigenvectors. Optional filters can be applied to remove edges based on edge length and direction. The output of this step is a graph representation of density ridges, without imposing prior assumption on its structure type and does not require all cells to be connected. To construct a second graph representation that connects every cell, we construct a minimum spanning tree graph with two additional steps: 1. Add edges to connect every connected component to its nearest neighbor in each of the other connected components. 2. Extract a minimum spanning tree of the whole graph. For use with dynbenchmark package, we further convert a graph to a dynbenchmark-compatible graph format with backbone cells assigned based on betweenness centralities. Cells that passed a threshold of 10 times the number of cells are assigned as backbones of the graph.

### Data and preprocessing

The 339-dataset benchmark dataset published by Saelens et al.^19^ was downloaded from https://zenodo.org/record/1443566. The unnormalized performance scores were extracted from https://github.com/dynverse/dynbenchmark_results/blob/1ac55e6c54a950890208b1f7730092d39783dfd2/06-benchmark/benchmark_results_unnormalised.rds. The normalized scores were computed as in ^19^, with the scaling factors kept to the same values as the original methods benchmarked. Other singe-cell datasets analyzed in this manuscript were from the following publications^11–13,15,20,24^. We created a Zenodo record for https://zenodo.org/record/3710980 that contains all the input data used in this manuscript.

---

**Supplementary Figure 1.**
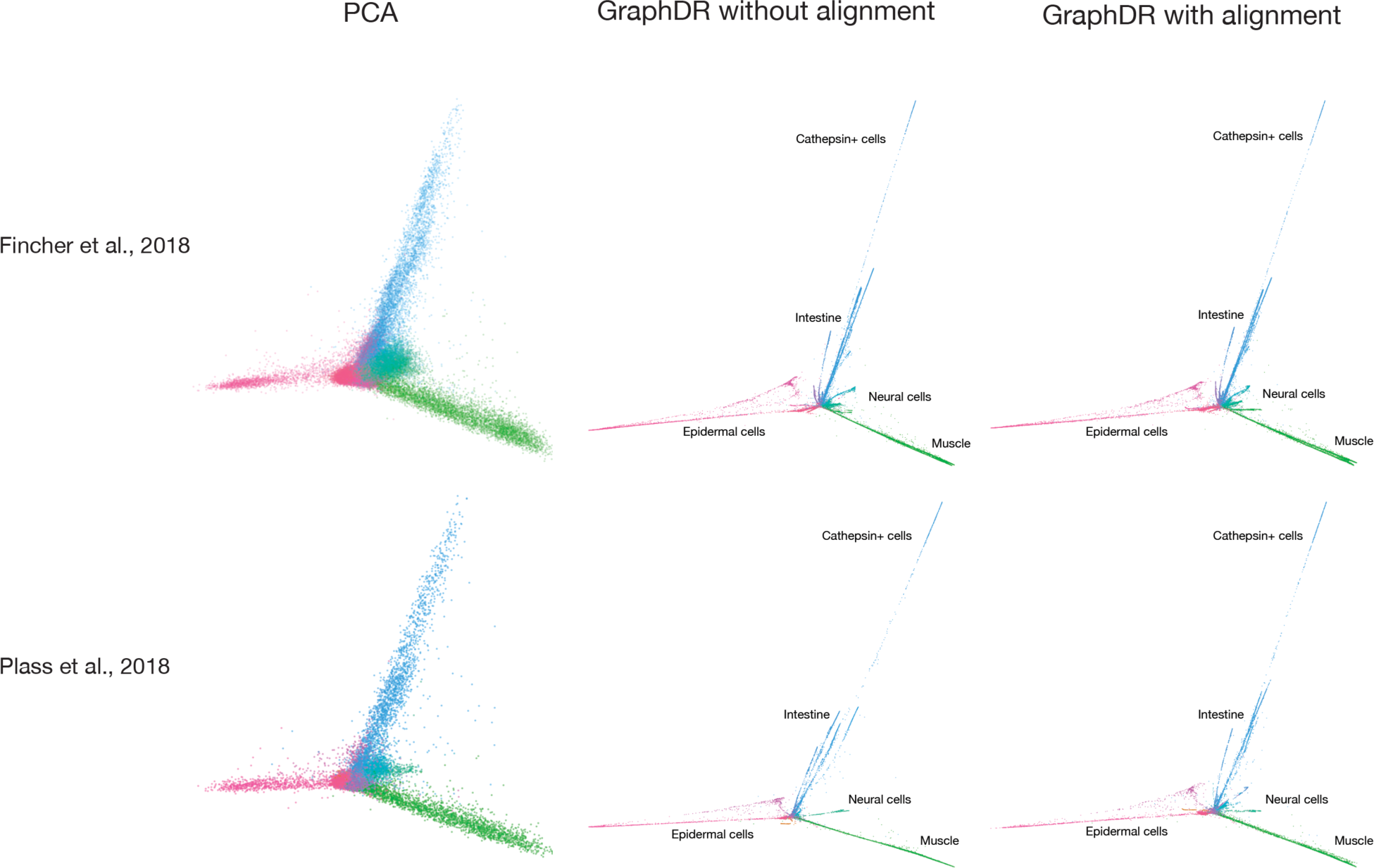
Dataset alignment with GraphDR further improves dataset comparison. GraphDR can be used to visualize and compare datasets while applied separately with a common linear subspace for quick comparison. By constructing graph jointly considering the experimental design (equivalent to batch design in **Supplementary Figure 2**), GraphDR effectively perform on-the-fly dataset alignment during the visualization and enables more precise comparison across datasets.

**Supplementary Figure 2.**
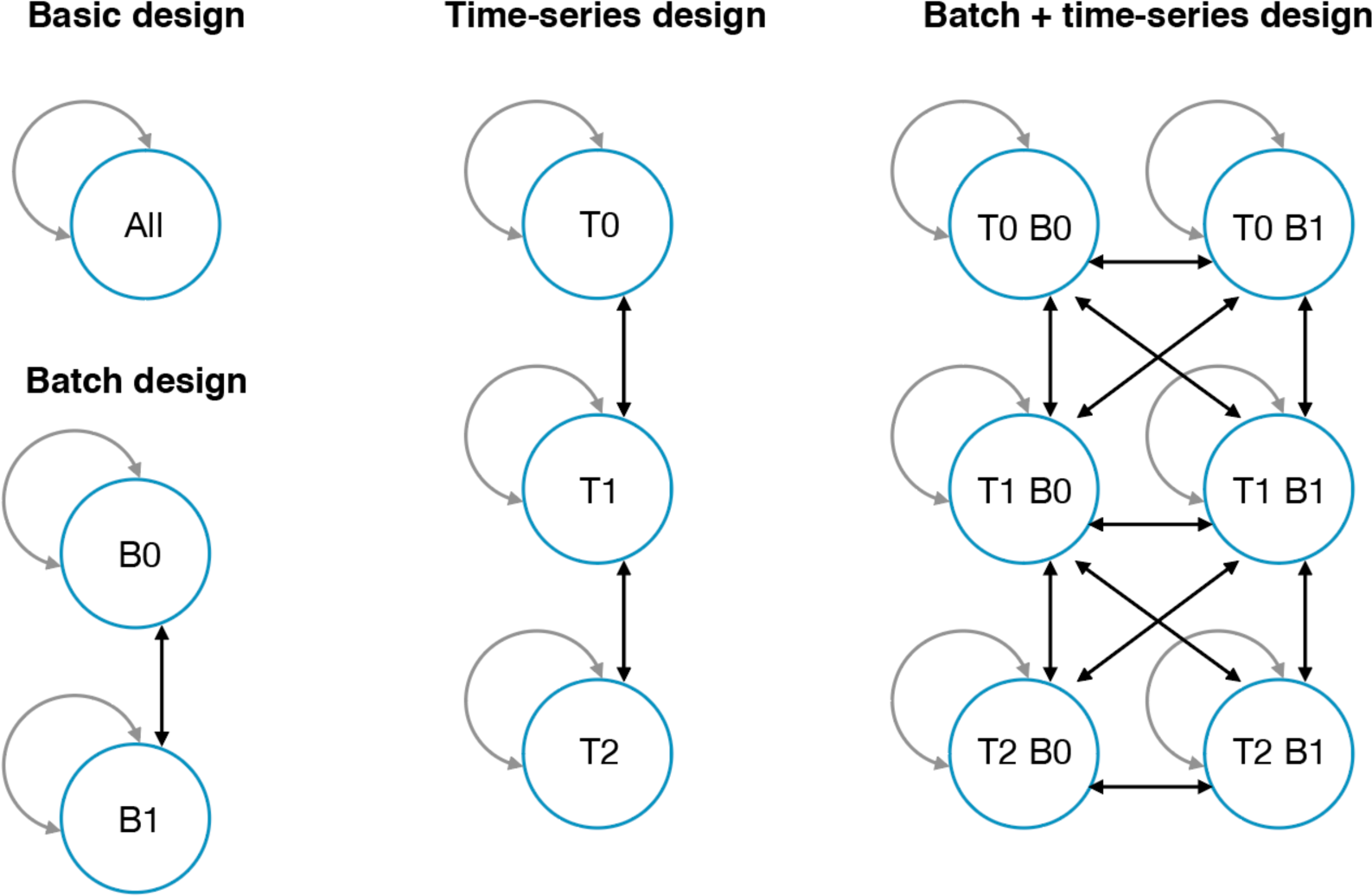
Experimental design encoding through graph construction. Experimental design information can be encoded through graph construction in GraphDR. Each arrow indicates that nearest-neighbor connections are established between the two groups, where two connected cells are in the two different groups. Self-loop indicates nearest-neighbor connections from cells within a group. Basic design constructs a nearest neighbor graph using all cells, which is suitable for single-batch experiments or experiments with minimal batch effects. Batch design addresses batch effects by introducing nearest-neighbor connections between all pairs of batches, in addition to with-in batch nearest-neighbor connections. Time-series design extends basic design by only allowing connections between the same and adjacent time points. Batch + time series design introduces nearest neighbor connections between two batches in the same or adjacent time points.

**Supplementary Figure 3.**
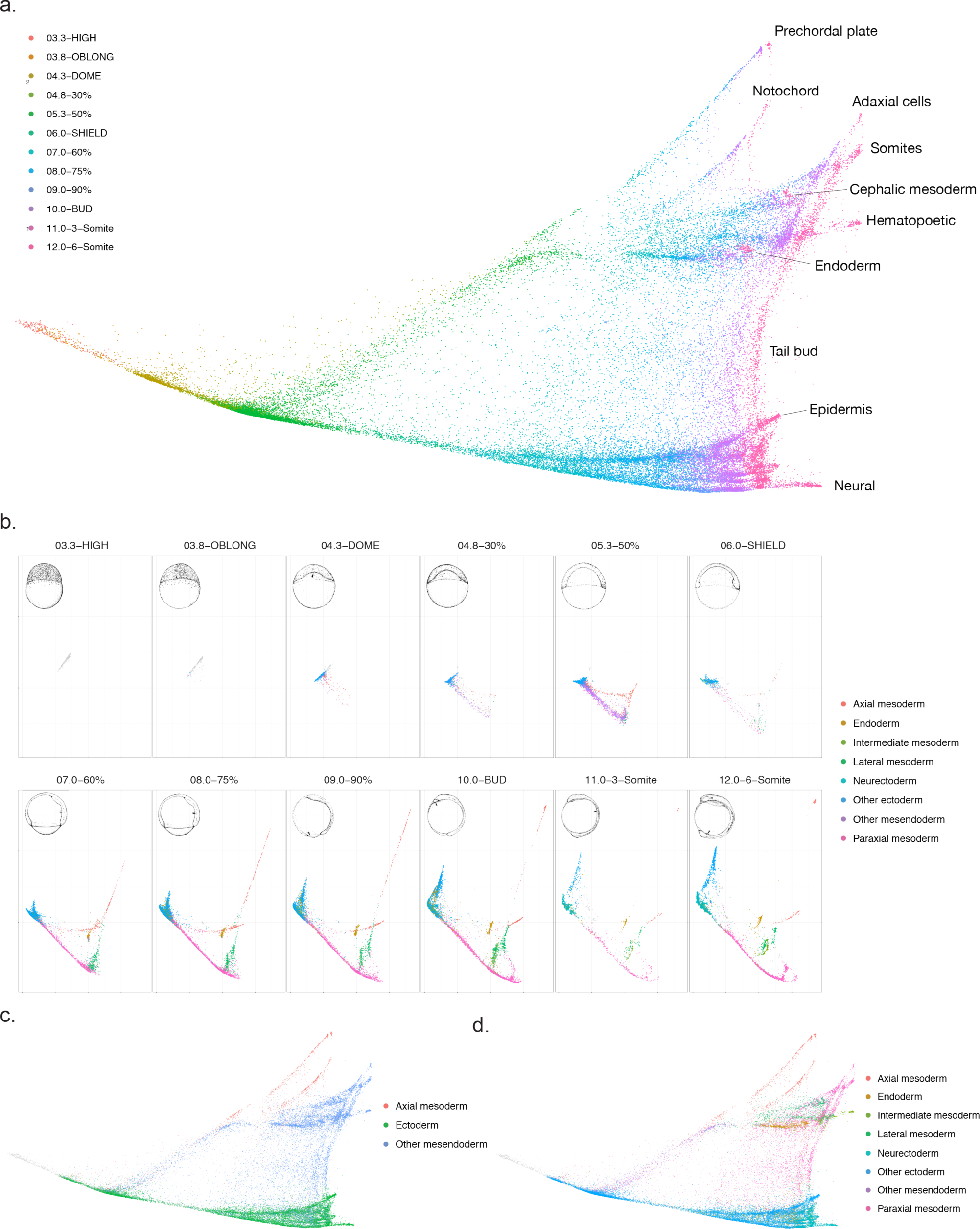
Visualization of Zebrafish whole embryo single-cell developmental landscape with GraphDR. Application of GraphDR to a single-cell dataset^15^ with a time-series design. a. Single-cell visualization by GraphDR, colored by developmental stages. b. Comparative visualization of developmental stages. This shows the “cross-section” view by visualizing the second and third dimensions. c-d. Single-cell visualization by GraphDR, colored by cell origins.

**Supplementary Figure 4.**
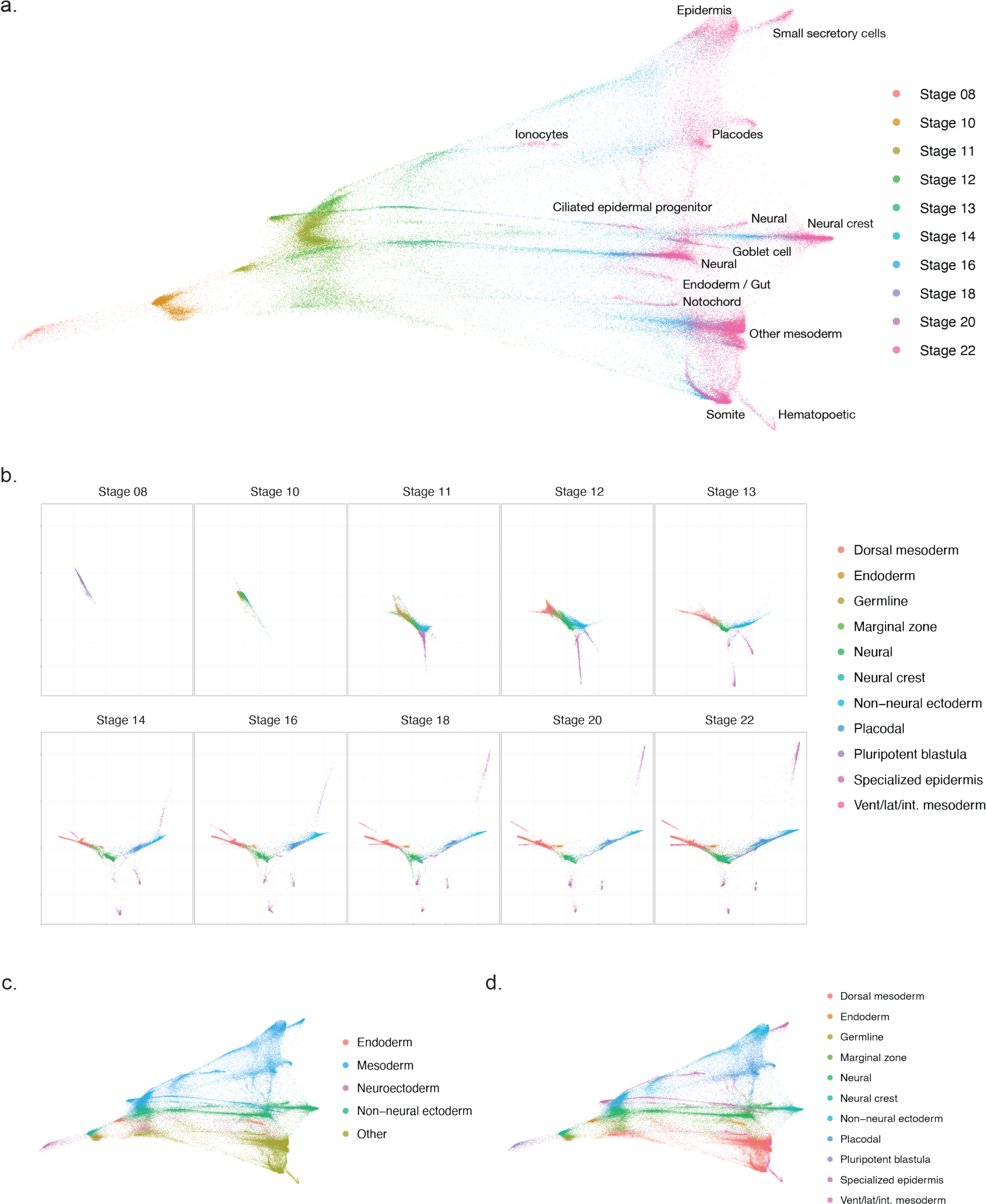
Visualization of Xenopus tropicalis whole embryo single-cell developmental landscape with GraphDR. This is an example of applying GraphDR to a single-cell dataset with a batch+time-series design. a. Single-cell visualization by GraphDR, colored by developmental stages. b. Comparative visualization of developmental stages. This shows the “cross-section” view by visualizing the second and third dimensions. c-d. Single-cell visualization by GraphDR, colored by cell origins.

**Supplementary Figure 5.**
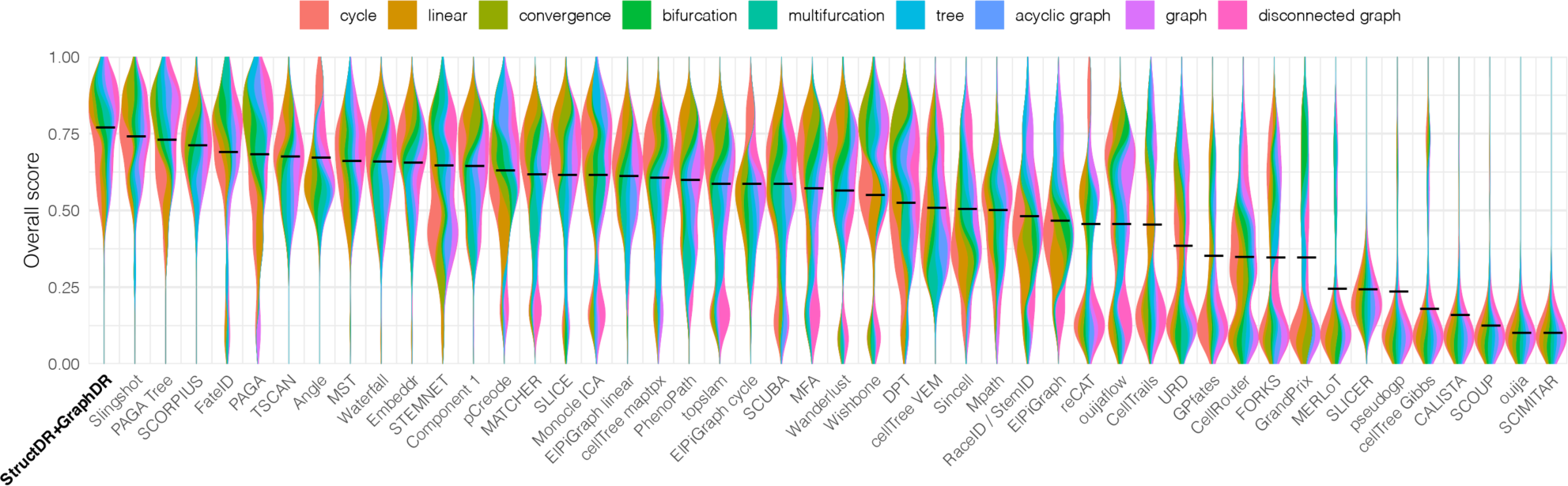
Performance score distributions on 339 dataset benchmark shown by dataset type. Per-dataset performance scores are computed based on Saelens et al. 2019^19^. The performance score distributions are shown with violin plots, broken down by dataset types.

**Supplementary Figure 6.**
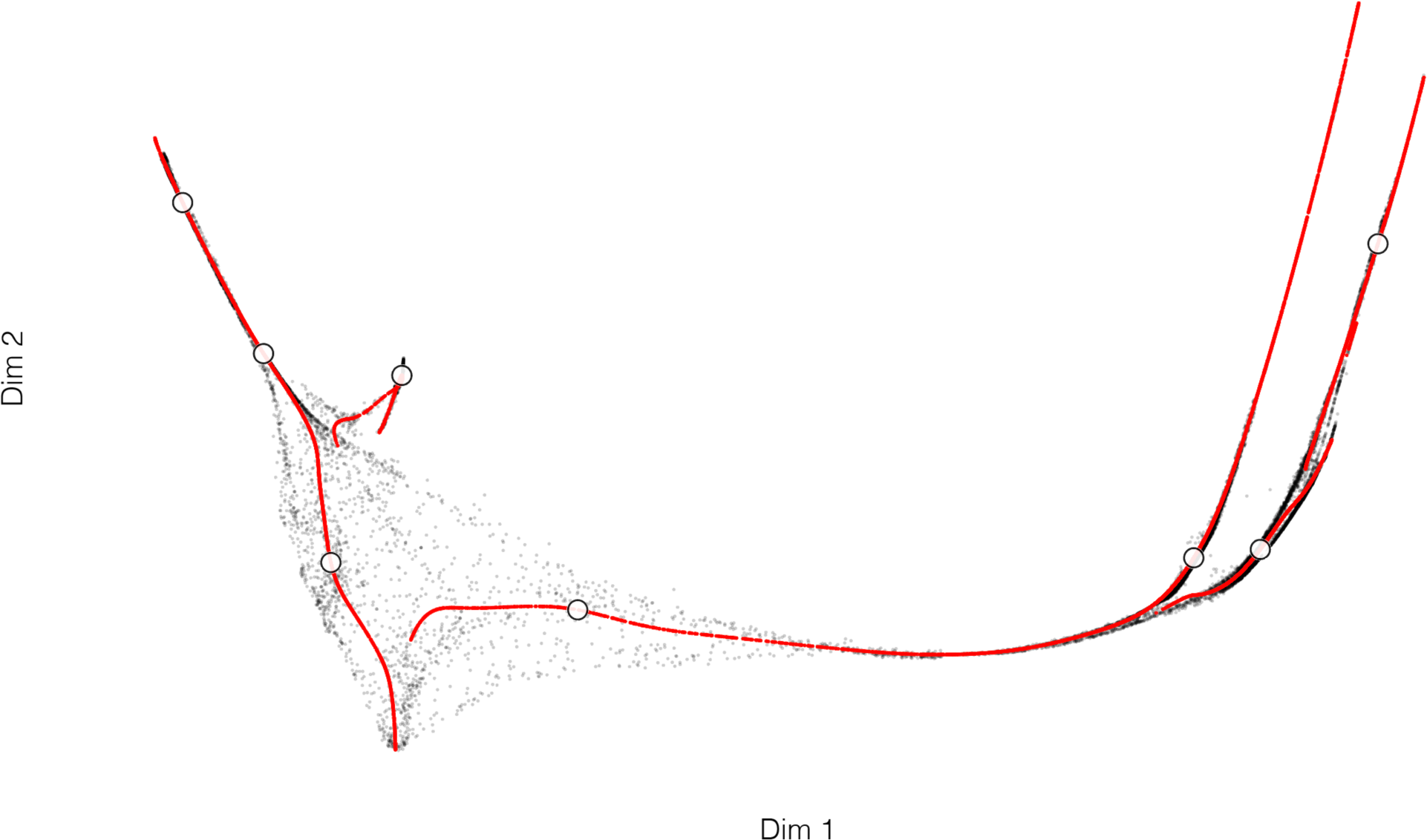
Trajectory identification with zero, one, and two dimensional density ridges example on a developmental hippocampus single-cell dataset. The circle symbols indicate zero-dimensional density ridge positions (local maxima of density function). The red dots indicate one-dimensional density ridge positions (trajectory). The black dots indicate two-dimensional density ridge positions.

**Supplementary Figure 7.**
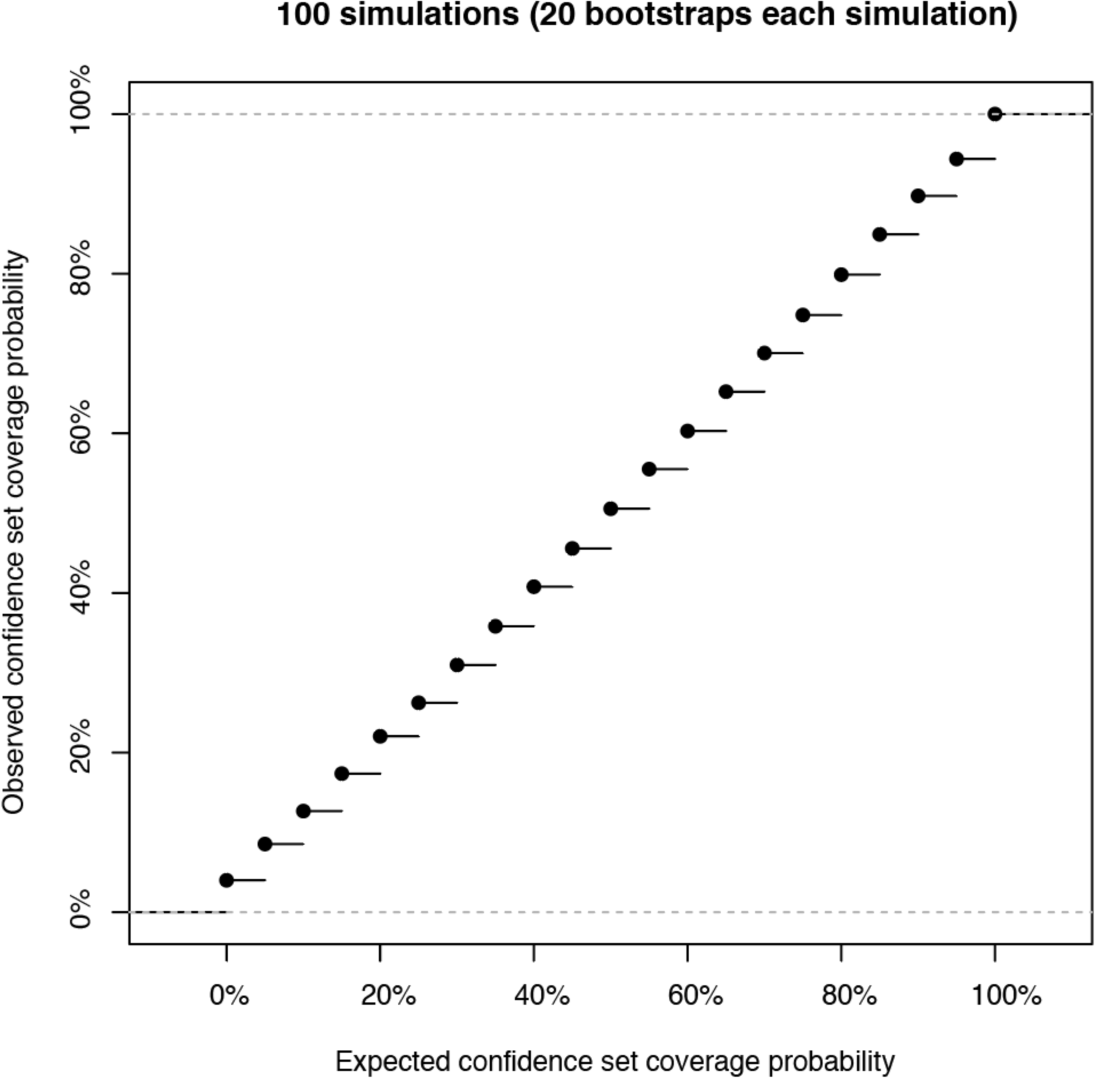
Simulation studies of confidence sets construction with nonparametric ridge estimation. 100 simulation datasets were generated. For each dataset the trajectory and confidence sets of each estimated trajectory were estimated with 20 bootstraps. x-axis shows the expected coverage probabilities of the constructed confidence sets. y-axis shows the observed proportion that the true trajectory position is covered by the confidence set.

## Code availability

All methods described in this manuscript are implemented in an open-source python package quasildr https://github.com/jzthree/quasildr.

